# Disruption of the microphysiological niche alters matrix deposition and causes loss of mitochondrial Ca^2+^ control in skeletal muscle fibers

**DOI:** 10.1101/2021.03.29.437546

**Authors:** Charlotte Gineste, Sonia Youhanna, Sabine U. Vorrink, Sara Henriksson, Andrés Hernández, Arthur J. Cheng, Thomas Chaillou, Andreas Buttgereit, Dominik Schneidereit, Oliver Friedrich, Kjell Hultenby, Joseph D. Bruton, Niklas Ivarsson, Linda Sandblad, Volker M. Lauschke, Håkan Westerblad

## Abstract

Cells rapidly lose their physiological phenotype upon disruption of their extracellular matrix (ECM)-intracellular cytoskeleton interactions. Here, we investigated acute effects of ECM disruption on cellular and mitochondrial morphology, transcriptomic signatures, and Ca^2+^ handling in adult mouse skeletal muscle fibers. Adult skeletal muscle fibers were isolated from mouse toe muscle either by collagenase-induced dissociation of the ECM or by mechanical dissection that leaves the contiguous ECM intact. Experiments were generally performed four hours after cell isolation. At this time, there were striking differences in the gene expression patterns between fibers isolated with the two methods; 24h after cell isolation, enzymatically dissociated fibers had transcriptomic signatures resembling dystrophic phenotypes. Mitochondrial appearance was grossly similar in the two groups, but 3D electron microscopy revealed shorter and less branched mitochondria in enzymatically dissociated than in mechanically dissected fibers. Similar increases in free cytosolic [Ca^2+^] during repeated tetanic stimulation were accompanied by marked mitochondrial Ca^2+^ uptake only in enzymatically dissociated muscle fibers. The aberrant mitochondrial Ca^2+^ uptake was partially prevented by the mitochondrial Ca^2+^ uniporter inhibitor Ru360 and by cyclosporine A and NV556, which inhibit the mitochondrial protein Ppif (also called cyclophilin D). Importantly, inhibition of Ppif with NV556 significantly improved survival of mice with mitochondrial myopathy in which muscle mitochondria take up excessive amounts of Ca^2+^ even with an intact ECM. In conclusion, skeletal muscle fibers isolated by collagenase-induced dissociation of the ECM display aberrant mitochondrial Ca^2+^ uptake, which involves a Ppif-dependent mitochondrial Ca^2+^ influx resembling that observed in mitochondrial myopathies.

## Main

The intracellular secondary messenger Ca^2+^ is of critical importance for a plethora of cellular functions. Within the cell, Ca^2+^ levels are compartmentalized and can differ by orders of magnitude between the cytosol and organelles, such as the sarcoplasmic reticulum (SR) and mitochondria. Dysregulation of intracellular Ca^2+^ handling is associated with a range of human pathologies, including diabetes [1, 2], neurodegenerative disease [3], cardiac failure [4], and chronic inflammatory conditions [5, 6]. In this context, mitochondrial Ca^2+^ levels have received increasing attention in recent years for their important roles in energy homeostasis, oxidative stress, and apoptosis [7, 8]. A limited and transient increase in free mitochondrial matrix [Ca^2+^] ([Ca^2+^]_mit_) can stimulate mitochondrial respiration and hence play an integral role in the regulation of cellular metabolism, whereas prolonged and excessive uptake can activate apoptotic and necrotic cell signaling pathways [9–11]. Tight control of [Ca^2+^]_mit_ seems particularly important in skeletal and cardiac muscle cells where cytosolic free [Ca^2+^] ([Ca^2+^]_cyt_) can reach µM concentrations, and even higher in the vicinity of SR Ca^2+^ release sites [12].

The mitochondrial Ca^2+^ uniporter (Mcu) was recently shown to be the pore-forming unit of a mitochondrial Ca^2+^ uptake channel [13, 14]. Besides the Mcu itself, mitochondrial Ca^2+^ entry is fine-tuned by various regulators [8, 15], as well as other, as of yet unidentified uptake pathways [16, 17]. [Ca^2+^]_mit_ is furthermore regulated by Ca^2+^ extrusion, which in skeletal muscle is mediated by the Na^+^/Ca^2+^ antiporter (Nclx)) [18].

Importantly, bidirectional mechanical and biochemical signaling between the extracellular matrix (ECM) and the intracellular cytoskeleton plays a key role in the control of cell structure, including the morphology and function of mitochondria [19–21]. In skeletal muscle fibers, mitochondria are tethered to the SR at sites of Ca^2+^ release [22], and this spatial organization is required for appropriate Ca^2+^ handling [23]. Disturbance of the spatial interaction between mitochondria and the SR in muscle cells can result in altered Ca^2+^ exchange between the two organelles, which in turn can entail functional impairments in both organelles [24]. For instance, disruption of crosstalk between the ECM and cytoskeleton due to mutations in ECM components give rise to several muscular dystrophies [25, 26], including collagen VI mutations resulting in Bethlem myopathy, Ullrich congenital muscular dystrophy, and congenital myosclerosis where mitochondrial Ca^2+^ overload is regarded as a key pathogenic factor [27]. Furthermore, muscle defects in collagen VI-deficient mice as well as in patients with collagen VI mutation-dependent muscular dystrophies are mitigated by pharmacological inhibition of the mitochondrial matrix protein peptidyl-prolyl cis-trans isomerase F (Ppif, NP_598845; also called cyclophilin D), a treatment that is considered to inhibit the Ca^2+^-dependent opening of the mitochondrial permeability transition pore [28–31].

In mammals, each muscle fiber is under the direct control of a single branch of an α-motoneuron and hence its cellular activation occurs independently of the activity of neighboring muscle fibers. Thus, studies on isolated single muscle fibers will reflect the function of muscle fibers *in vivo* and are therefore highly valuable in mechanistic studies of, for instance, the intracellular signaling associated with muscle activation and contraction. Importantly, single muscle fibers are often isolated via collagenase treatment [23, 32–36], which inevitably disrupts the extracellular niche and ECM-cell interactions. Here, we compare collagenase-dissociated mouse flexor digitorum brevis (FDB) fibers with fibers obtained by high-precision mechanical dissection [37] and demonstrate that an intact microenvironment is required for maintenance of cellular structure and mitochondrial Ca^2+^ handling. Specifically, by integrating data obtained with Second Harmonic Generation (SHG) imaging, electron microscopy and RNA-sequencing, we show that enzymatic fiber dissociation, but not mechanical microdissection, causes a rapid breakdown of cellular organization and the deterioration of molecular signatures characteristic of mature muscle fibers. Furthermore, Ca^2+^ imaging and functional assays in healthy and mitochondrial myopathy fibers show that only microdissected fibers retain their ability to control mitochondrial Ca^2+^ influx during contraction-associated increases in [Ca_2+_]_cyt_.

## Results

### Intact links between ECM and cytoskeleton are necessary to maintain the structural integrity of skeletal muscle fibers

To examine the effects of the extracellular microenvironment on cellular phenotype, we compared mechanically dissected mouse FDB fibers with fibers isolated by conventional enzymatic dissociation using collagenase. Mechanical microdissection permits the isolation of muscle fibers with tendons attached and intact sarcolemma [37], including the adjacent ECM scaffold with preserved focal adhesion complexes. In contrast, physiological ECM contacts are destroyed in enzymatically isolated fibers. We first examined whether the different isolation methods translated into differences in cell morphology. To this end, we combined multi-photon-based Second Harmonic Generation (SHG) imaging and a quantitative morphometry technique to assess possible alterations in the myofibrillar architecture induced by the collagenase treatment [38]. SHG microscopy on single mechanically dissected FDB fibers revealed a well-defined complex myofibrillar architecture with longitudinal ridges and valleys. Enzymatically dissociated fibers, on the other hand, were more homogeneously delineated with almost circular cross-sections and shorter, more evenly extended sarcomeres (Figure 1A & B; Videos 1 & 2).

**Figure 1.**
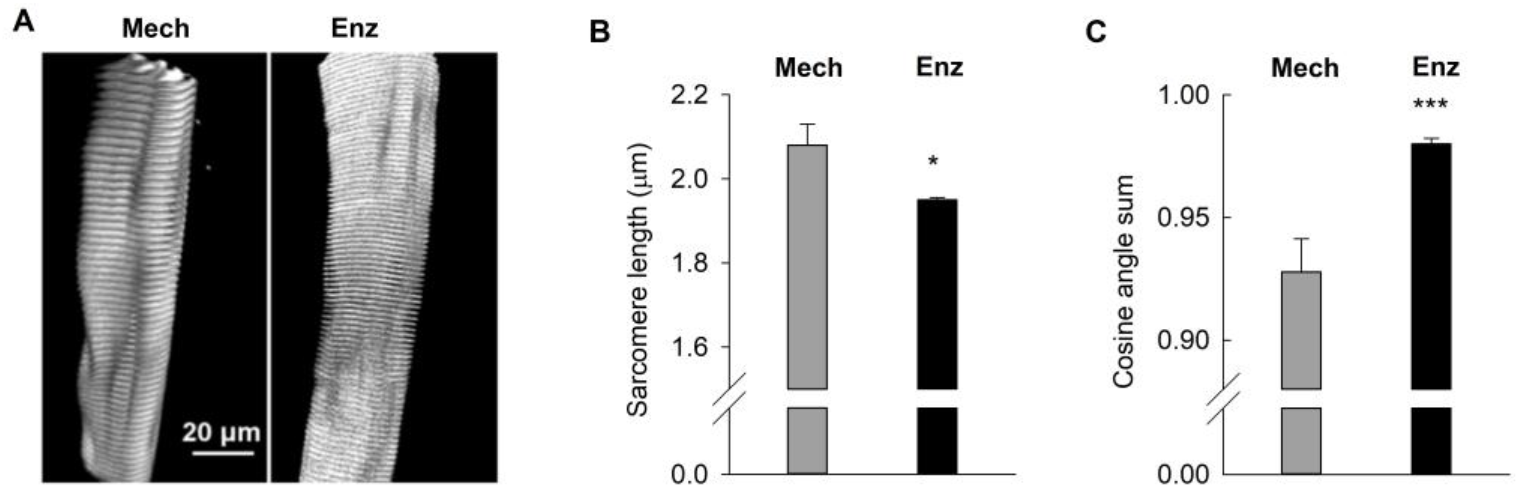
Disruption of the extracellular matrix results in altered cell shape and downregulation of genes encoding for important structural proteins. (**A**) Representative SHG images from a mechanically dissected (Mech) and an enzymatically dissociated (Enz; using collagenase) mouse FDB fiber. Scale bar, 20 µm. Average data of (**B**) sarcomere length and (**C**) cosine angle sum (CAS) in mechanically dissected (n = 5) and enzymatically dissociated (n = 13) fibers. CAS of 1 represents a perfectly linear parallel pattern, whereas CAS = 0 reflects perpendicularly oriented structures. Grey bars = mechanically dissected fibers; black bars = enzymatically dissociated fibers; experiments were performed 4 hours after muscle fiber isolation. * P < 0.05 and *** P < 0.001 vs. mechanically dissected fibers with unpaired t-test. Data are presented as mean ± SEM.

To further characterize the structural difference between mechanically dissected and enzymatically dissociated fibers, we quantified the cosine angle sum (CAS) in SHG images as a measure of a fiber’s myofibrillar angular alignment [38, 39]. Notably, CAS was significantly higher (reflecting a more linear parallel pattern) for enzymatically dissociated than for mechanically dissected fibers, further corroborating the observation of homogeneously striated patterns and parallel myofibrillar alignment in dissociated fibers (Figure 1C).

### 3D electron microscopy revealed an enzymatic dissociation-induced reduction in volume of individual mitochondria

Next, we evaluated whether the collagenase-induced disruption of the extracellular microenvironment would propagate into alterations in mitochondrial spatial organization and ultrastructure. To this end, we analyzed fibers of both mechanically dissected and enzymatically dissociated fibers using transmission electron microscopy (TEM) and focused ion beam scanning electron microscopy (FIB-SEM) (Figure 2). Notably, TEM analysis of ultrathin sections showed no significant difference in mitochondrial cross-sectional areas between the two groups (Figure 2C), whereas mechanically dissected fibers featured longer and more branched mitochondria when assessed with FIB-SEM [40, 41]. Thus, by sequential FIB milling of 30 nm slices, 3D mitochondrial models of similar fiber volumes (∼365 µm^3^) were obtained from a mechanically dissected and an enzymatically dissociated fiber (Figure 2D-G; Videos 3 & 4). The 3D images look grossly similar in the two fibers and the total mitochondrial volume was similar in the two models (20.2 *vs*. 17.1 µm^3^, corresponding to 5.5% *vs*. 4.6% of the total fiber volume). On the other hand, the number of mitochondria was clearly lower (86 *vs*. 140) and their average volume larger in the mechanically dissected (0.235 µm^3^) than in the enzymatically dissociated (0.122 µm^3^) fiber.

**Figure 2.**
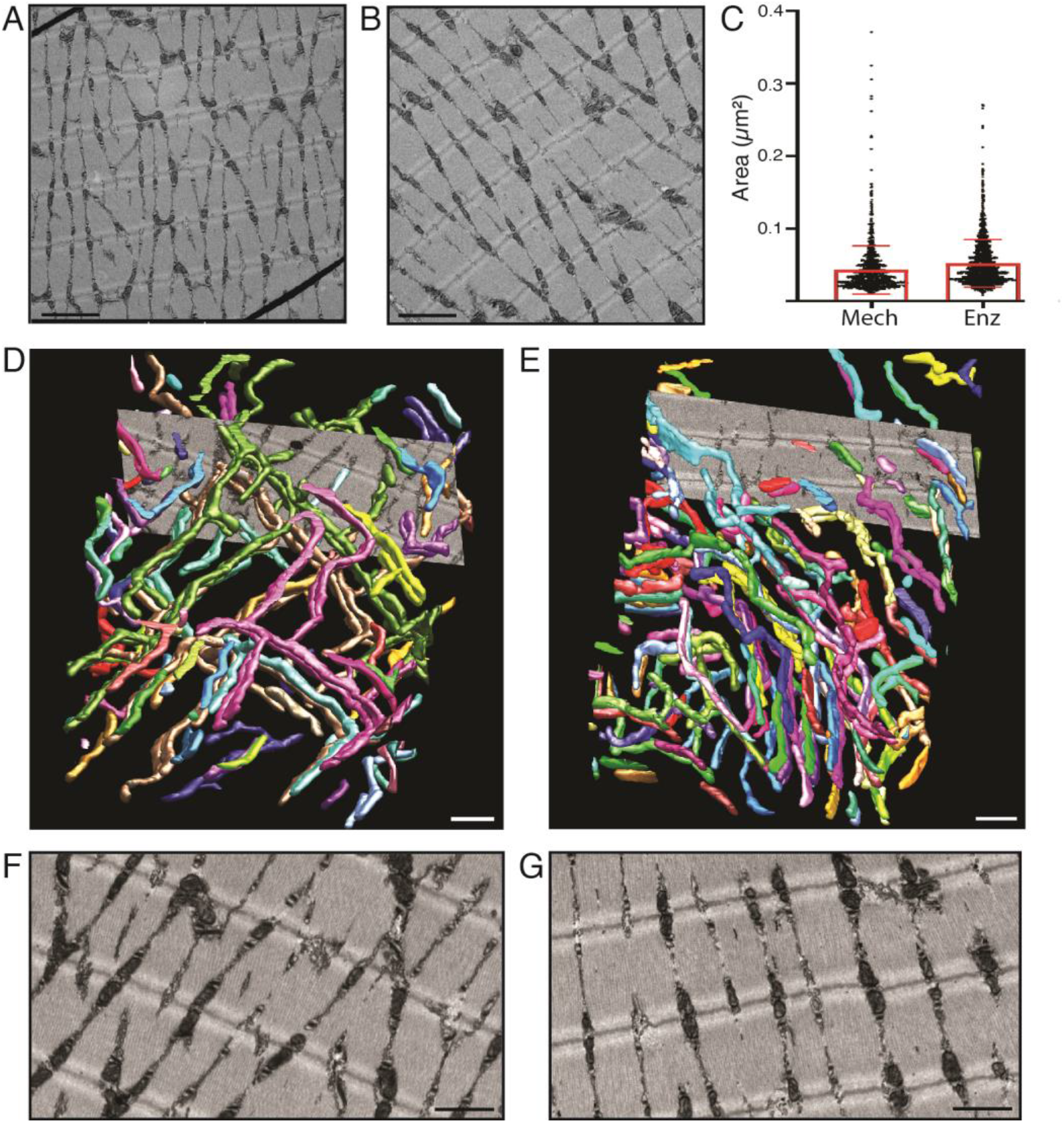
Disruption of the extracellular matrix causes subtle changes in mitochondrial morphology as detected with 3D electron microscopy. Representative TEM images of a mechanically dissected (**A**) and a collagenase dissociated (**B**) FDB muscle fiber. Scale bar in A and B is 2 µm. (**C**) Graphical presentation of individual and mean (± SD) mitochondrial areas measured from TEM images acquired from mechanically dissected fibers (left; n=1154 mitochondria in 20 images of distinct areas of 6 fibers) and enzymatically dissociated fibers (right, n=1390 mitochondria in 20 images of distinct areas of 10 fibers). 3D model of the mitochondria in a volume of 365 µm^3^ muscle tissue obtained with FIB-SEM in a dissected fiber (**D**) and collagenase-treated fiber (**E**); rotating 3D models are presented in Videos 3 & 4. Individual mitochondria are distinguished by different colors. Representative images from slices used in the FIB-SEM-based modelling acquired in the dissected (**F**) and the collagenase-treated (**G**) fiber, respectively. Fibers were fixed 4 hours after being isolated. Scale bar in D-G is 1 µm.

### Disruption of the cellular microenvironment causes a rapid deterioration of transcriptomic signatures

We used RNA-sequencing to compare the transcriptomic signatures of freshly isolated muscle fiber bundles to fibers isolated either by microdissection or enzymatic dissociation. Notably, we observed striking differences in gene expression patterns depending on the isolation method (Figure 3A). Principal component analysis revealed that overall changes in expression were more rapid and extensive in dissociated fibers (Figure 3B). Compared to freshly isolated muscle fiber bundles, 514 genes were differentially expressed in microdissected fibers, whereas more than twice as many (1156 genes) were altered upon dissociation (FDR=0.01). Systematic comparison of expression differences between isolation methods revealed that after 4h, 3,210 genes showed significantly higher levels in microdissected than in enzymatically dissociated fibers and these were enriched in complement signaling, ECM proteoglycans, ephrin signaling and rRNA processing (Figure 3C). In contrast, only 85 genes showed higher expression in dissociated than in microdissected fibers, specifically those involved in glycogen synthesis and gluconeogenesis. Conversely, 24h after isolation, microdissected fibers featured 1,203 significantly upregulated genes mostly related to ECM biogenesis and organization, whereas enzymatically dissociated fibers show upregulation of 2,323 genes significantly enriched in cell cycle, DNA replication and repair (Figure 3D).

**Figure 3.**
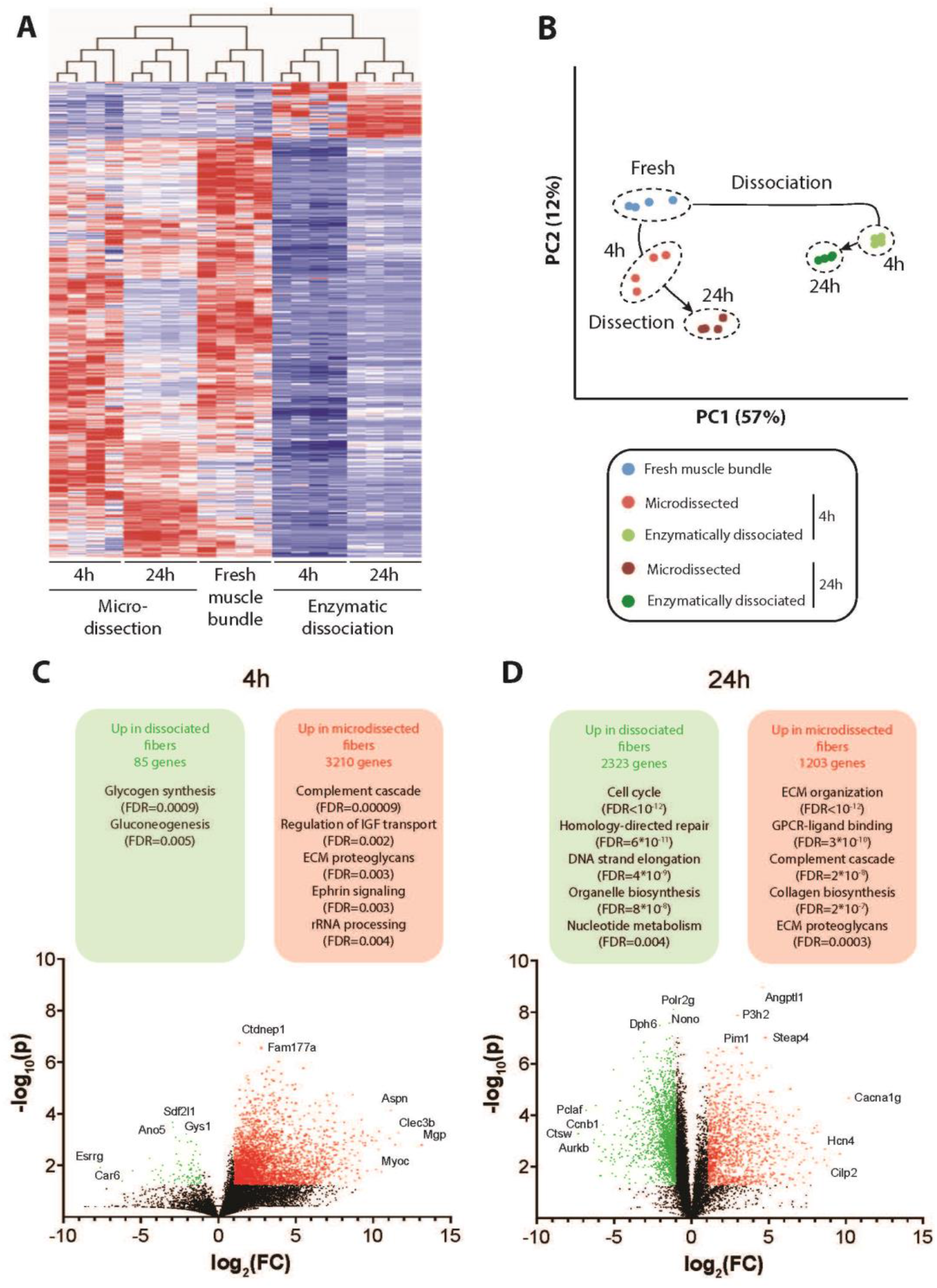
RNA-Sequencing reveals major transcriptomic signature changes in response to muscle fiber isolation. (**A)** Mean-centered, sigma-normalized heatmap visualization of differentially expressed genes (F-test; FDR=5%; Benjamini-Hochberg correction). Note that microdissected fibers more closely resemble the gene expression patterns of freshly isolated muscle bundles than enzymatically dissociated fibers. (**B)** The corresponding principal component analysis (PCA) reveals that enzymatic dissociation and microdissection result in orthogonal gene expression alterations. (**C-D)** Volcano plots visualizing differentially expressed genes between microdissected and enzymatically dissociated fibers 4h (**C**) and 24h (**D**) after isolation. Red and green dots indicate genes that are significantly (fold-change [FC]>2 and p<0.05 in heteroscedastic two-tailed T-test) upregulated in microdissected and enzymatically dissociated fibers, respectively. Significantly enriched pathways corresponding to the differentially expressed genes are indicated in the respective colored boxes.

Notably, the expression of genes encoding for myosin heavy chain IIX (Myh1), IIB (Myh4), and I (Myh7) was substantially reduced after 4h irrespective of isolation method; however, after 24h expression had recovered and was significantly higher in microdissected than in dissociated fibers (Figure 4A), whereas the gene encoding for myosin heavy chain IIA (Myh2) was not differentially expressed between the two groups (Supplementary Figure 1A).

**Figure 4.**
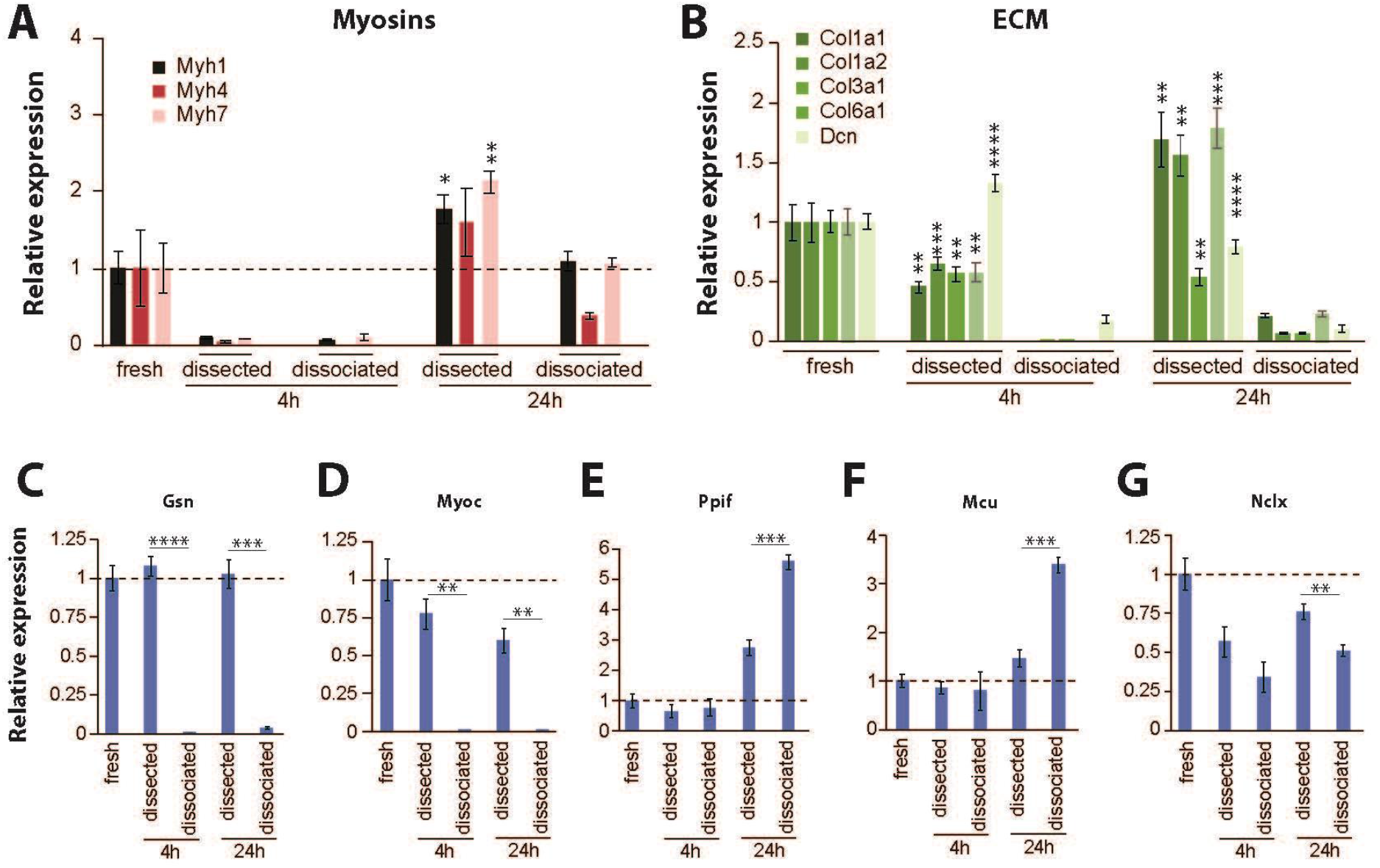
Enzymatic dissociation alters the expression of myosins, extracellular matrix components and mitochondrial Ca^2+^ regulators. Expression of functionally relevant myosins (**A**) and extracellular matrix (ECM) genes (**B**) are significantly downregulated in dissociated vs. dissected fibers. The actin regulators Gsn (**C**) and Myoc (**D**) are among the most affected genes whose expression is almost completely lost upon enzymatic dissociation. The gene expression levels of mitochondrial proteins involved in Ca^2+^ uptake, Ppif (**E**) and Mcu (**F)**, are increased in dissociated fibers, whereas levels of Nclx involved in mitochondrial Ca^2+^ efflux are significantly decreased (**G**). *, **, *** and **** indicate P < 0.05, P < 0.01, P < 0.001 and P < 0.0001 between dissected and dissociated fibers at the same time point. Data are presented as mean ± SEM; n=4.

The gene expression of critical ECM components, including the abundant muscle collagens Col1a1, Col1a2, Col3a1, and Col6a1, as well as the proteoglycan decorin (Dcn), were almost entirely lost in dissociated fibers at both time points, whereas expression remained relatively stable in microdissected fibers (Figure 4B). Similar effects were observed for the gene expression of gelsolin (Gsn), a calcium-regulated protein involved in the assembly of actin filaments, and myocilin (Myoc), a regulator of cytoskeletal function (Figure 4C-D). Interestingly, at 24h the expression of *Ppif* and *Mcu* were higher with enzymatic dissociation than with microdissection (Figure 4E-F), whereas the expression of *Nclx* was lower in dissociated fibers (Figure 4G), hence all three differences acting towards increased mitochondrial Ca^2+^ accumulation in enzymatically dissociated fibers. The expression of MCU regulators showed no consistent pattern (Supplementary Figure 1B-F). Taken together, these findings confirm that enzymatic dissociation of muscle fibers results in a rapid deterioration of the mature muscle phenotype, approaching signatures that resemble dystrophic phenotypes with decreased expression of genes encoding for critical ECM elements and ECM-cytoskeletal signal transducers, as well as an activation of regenerative pathways.

### Collagenase treatment results in aberrant mitochondrial Ca^2+^ uptake

Next, we investigated whether collagenase-induced disturbance of the ECM would impact cellular Ca^2+^ handling. We previously showed that repeated tetani do not result in markedly increased [Ca^2+^]_mit_ in mechanically dissected FDB fibers of wildtype mice, implying intact mitochondrial Ca^2+^ control [42]. Intriguingly and in sharp contrast to mechanically dissected fibers, we noticed a marked increase in [Ca^2+^]_mit_, as indicated by an increased fluorescence signal of the mitochondrial Ca^2+^ indicator rhod-2, when FDB fibers were isolated by enzymatic digestion (Figure 5A). Thus, after enzymatic dissociation, rhod-2 fluorescence was more than doubled immediately after 25 brief (350 ms duration) tetani (Figure 5B). In contrast, after mechanical dissection, [Ca^2+^]_mit_ was only marginally increased after repeated tetani, irrespective of whether fibers contracted isometrically or were allowed to shorten freely. However, collagenase treatment of mechanically dissected fibers abolished these differences, demonstrating that the observed effects are not due to isolation but due to the intact collagen-containing microenvironment of the isolated fibers. Importantly, the differences in mitochondrial Ca^2+^ uptake between mechanically dissected and enzymatically dissociated fibers were not due to differences in tetanic [Ca^2+^]_cyt_ (Figure 5C), which indicates that defective mitochondrial Ca^2+^ control rather than general alterations in cellular Ca^2+^ handling underlie the aberrant elevation in [Ca^2+^]_mit._ during repeated contractions in enzymatically dissociated fibers.

**Figure 5.**
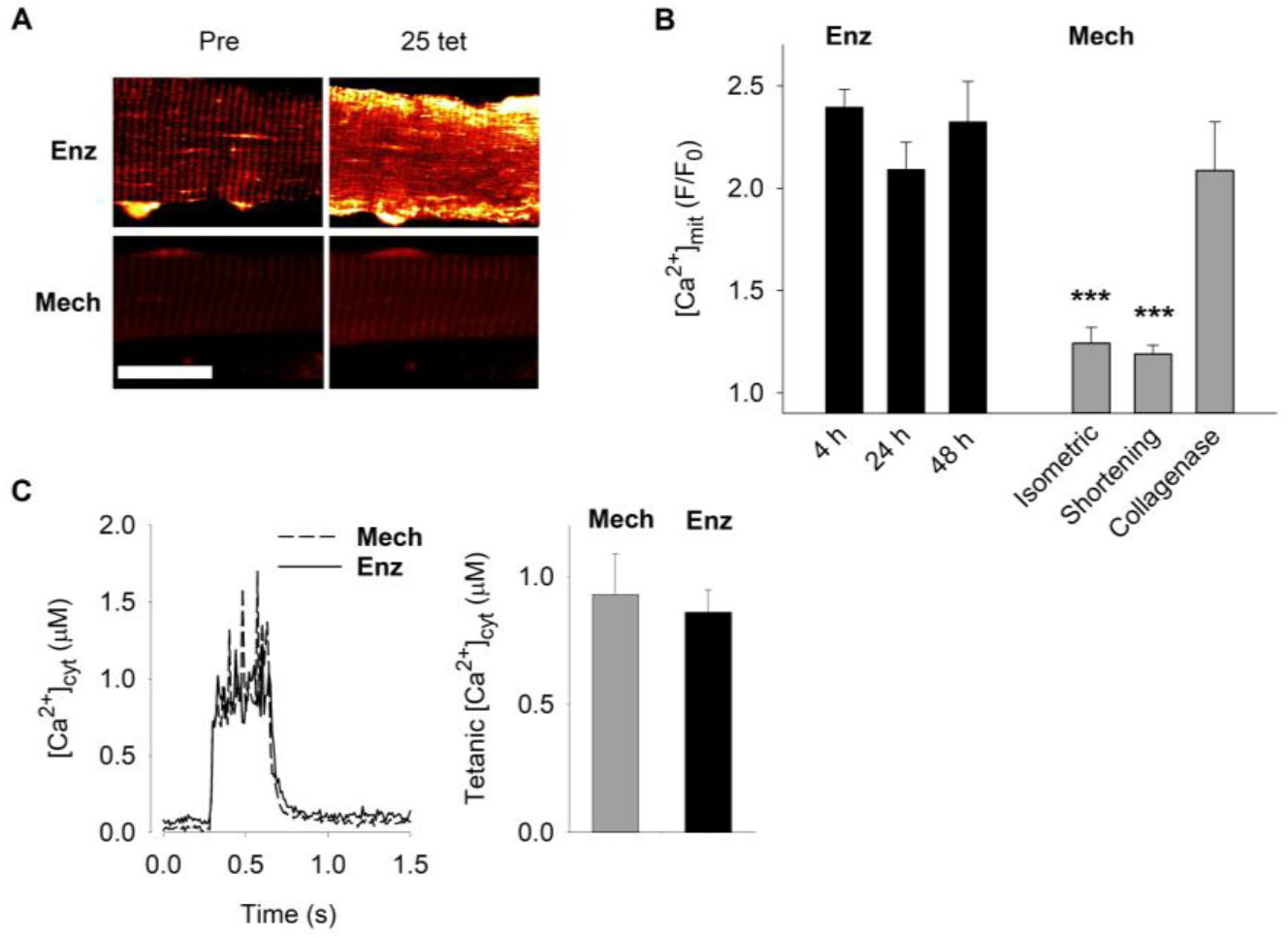
Enzymatic dissociation of muscle fibers results in impaired mitochondrial Ca^2+^ control. **(A)** Representative rhod-2 fluorescence images of enzymatically dissociated (Enz) and mechanically dissected (Mech) mouse FDB fibers before and after 25 brief tetanic contractions. Scale bar, 20 µm. **(B)** Mean data of rhod-2 fluorescence immediately after (F) relative to before (F_0_) 25 repeated tetani in enzymatically dissociated fibers stored for 4-48 hours after isolation (black bars) and in fibers studied 4 hours after mechanical dissection either contracting isometrically, shortening freely, or after collagenase treatment (grey bars). *** P < 0.001 vs. 4 hours dissociated fibers with one-way ANOVA (n ≥ 5). (**C**) Average [Ca^2+^]_cyt_ records (left) and mean tetanic [Ca^2+^]_cyt_ (right) during tetanic stimulation (70 Hz, 350 ms duration) of enzymatically dissociated (solid line and black bar) and mechanically dissected (dashed line and grey bar) FDB fibers (n = 6 in both groups). Data are presented as mean ± SEM.

### Mitochondrial Ca^2+^ uptake in enzymatically dissociated fibers is mediated via MCU- and PPIF-dependent pathways

To investigate the molecular mechanisms by which Ca^2+^ entered mitochondria in enzymatically dissociated cells, we used ruthenium red subcomponent Ru360 to inhibit MCU-mediated mitochondrial Ca^2+^ entry [43]. While Ru360 specifically inhibits MCU in experiments on isolated mitochondria, its limited plasma membrane permeability limits its use in intact cell systems [44]. Therefore, enzymatically dissociated FDB fibers were first microinjected with Ru360 and subsequently superfused by Ru360 for 30 minutes before commencing the repeated tetanic stimulation. Notably, Ru360 treatment significantly decreased but did not abrogate the tetanic stimulation-induced increase in [Ca^2+^]_mit_ (Figure 6A & B).

**Figure 6.**
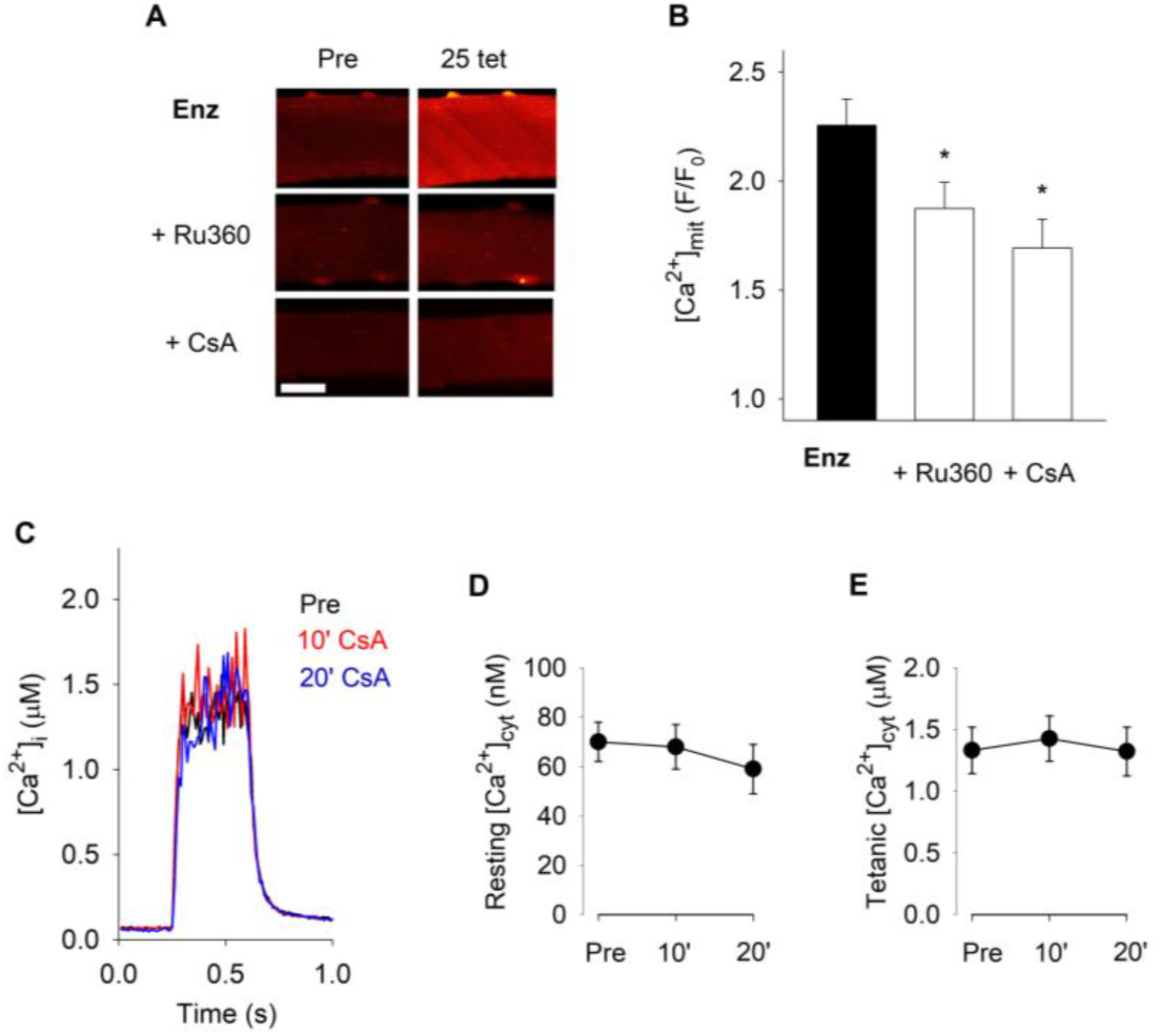
Aberrant mitochondrial Ca^2+^ uptake in enzymatically dissociated muscle fibers is mitigated by pharmacological inhibition of MCU and PPIF. (**A**) Representative rhod-2 fluorescence images of enzymatically dissociated FDB fibers obtained before (Pre) and after 25 brief tetanic contractions produced either under control conditions or in the presence of Ru360 (microinjected plus 10 µM superfusion) or cyclosporin A (CsA; 1.6 µM). Scale bar, 20 µm. (**B**) Mean data of the increase in rhod-2 fluorescence after (F) relative to before (F_0_) 25 repeated tetani; * P < 0.05 vs. control (Enz) with one-way ANOVA (n ≥ 13). (**C**) Superimposed average [Ca^2+^]_cyt_ records obtained in eight FDB fibers during tetanic stimulation (70 Hz, 350 ms duration) before (Pre) and after 10 and 20 min exposure to CsA (1.6 µM). Mean data of [Ca^2+^]_cyt_ at rest (**D**) and during the tetanic stimulation (**E**); one-way repeated measures ANOVA shows no effect of CsA exposure on either resting (P = 0.3) or tetanic (P = 0.5) [Ca^2+^]_cyt_ (n = 8). Experiments were performed 4 hours after fiber isolation. Data are presented as mean ± SEM.

Cyclosporin A (CsA) is a cyclic endecapeptide calcineurin inhibitor that binds to the mitochondrial protein PPIF (cyclophilin D) [45, 46] and counteracts the opening of the mitochondrial permeability transition pore [47–50].CsA has previously been shown to reduce mitochondrial Ca^2+^ uptake in mitochondrial myopathy muscle fibers exposed to repeated tetanic stimulation and in ischemic rabbit cardiomyocytes [42, 51, 52]. In enzymatically dissociated fibers exposed to 25 repeated tetani, CsA significantly blunted the increase in [Ca^2+^]_mit_ (Figure 6A & B), and the magnitude of reduction was similar to that observed with Ru360. Measurements of [Ca^2+^]_cyt_ showed no effect of CsA either at rest or during tetanic stimulation (Figure 6C-E).

These data suggest an important role of PPIF in mitochondrial Ca^2+^ control. To further investigate this possibility, we used the novel, specific cyclophilin inhibitor, NV556 [53]. Importantly, as with CsA, NV556 significantly lowered [Ca^2+^]_mit_ during repeated tetani as well as in the subsequent recovery period (Figure 7A & B). Moreover and in agreement with CsA, application of NV556 exposure had no effect on [Ca^2+^]_cyt_ at rest or during tetanic stimulation (Figure 7C-E). Thus, these results support a model in which the isolation of muscle fibers from their native microenvironment causes a dysregulation of cellular organization and defective control of Ca^2+^ influx into the mitochondrial matrix, which in part depends on MCU and PPIF.

**Figure 7.**
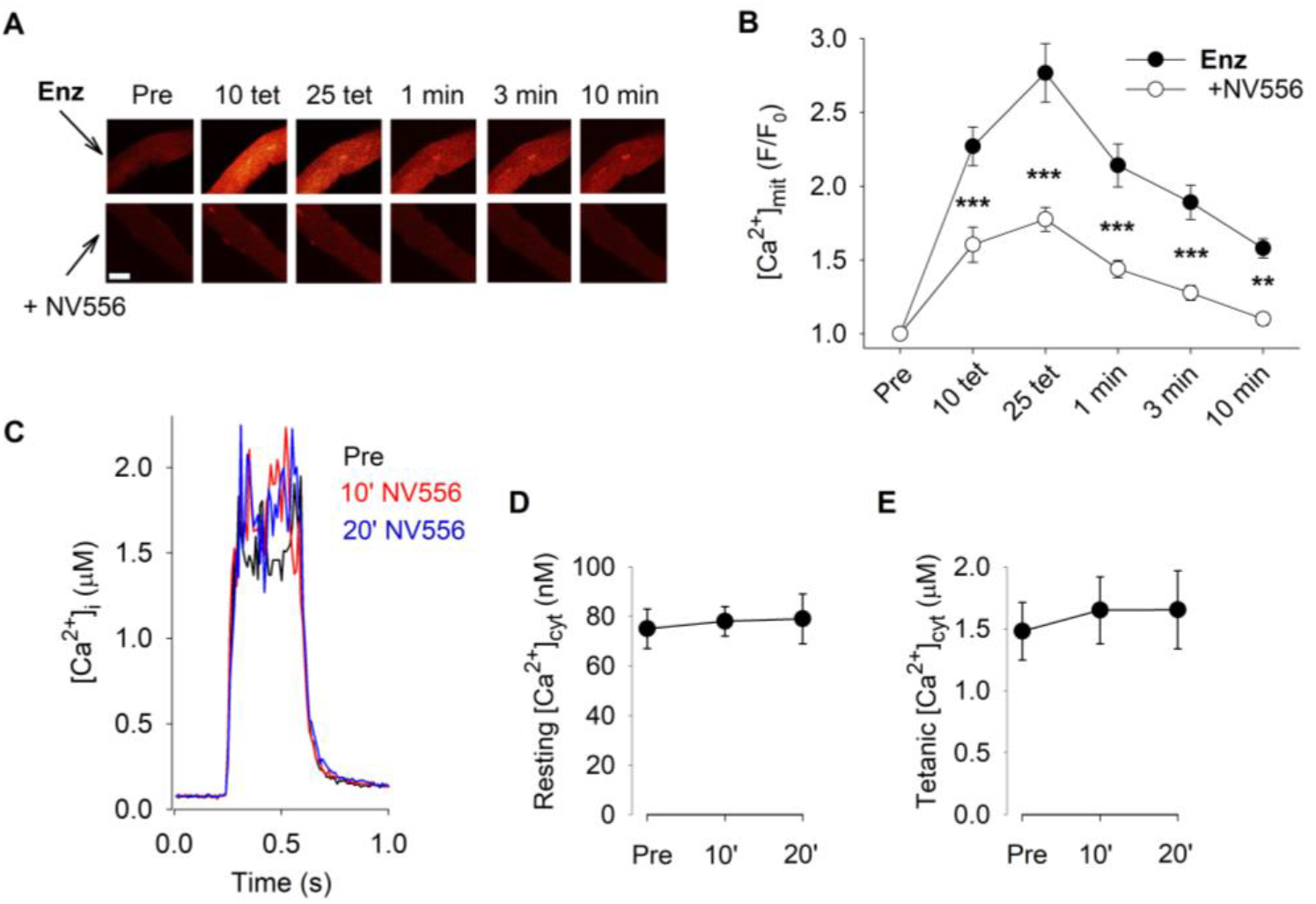
Aberrant mitochondrial Ca^2+^ uptake in enzymatically dissociated muscle fibers is mitigated by the cyclophilin inhibitor NV556. Representative rhod-2 fluorescence images (**A**) and quantification of the relative increase in fluorescence (**B**) of enzymatically dissociated FDB fibers stimulated with 25 repeated tetani either under control conditions or in the presence of the cyclophilin inhibitor NV556 (5 µM); ** P < 0.01, *** P < 0.001 vs. control with two-way repeated measures ANOVA (n ≥ 24). Scale bar, 20 µm. (**C**) Superimposed average [Ca^2+^]_cyt_ records obtained in nine FDB fibers during tetanic stimulation (70 Hz, 350 ms duration) before (Pre) and after 10 and 20 min exposure to NV556 (5 µM). Mean data of [Ca^2+^]_cyt_ at rest (**D**) and during the tetanic stimulation (**E**); one-way repeated measures ANOVA shows no effect of CsA exposure on either resting (P = 0.7) or tetanic (P = 0.5) [Ca^2+^]_cyt_ (n = 9). Experiments were performed 4 hours after fiber isolation. Data are presented as mean ± SEM.

Notably and consistent with previous reports [32], enzymatically dissociated fibers exposed to 25 repeated tetanic contractions did not display any marked depolarization of the mitochondrial membrane potential (Δ*ψ*_m_; measured with the fluorescent indicator TMRE) or any increase in reactive oxygen species (ROS; measured with the fluorescent indicator MitoSOX Red) (Supplementary Figure 2). These findings imply that the observed increases in [Ca^2+^]_mit_ upon enzymatic dissociation did not reach levels high enough to acutely trigger severe mitochondrial dysfunction.

### Enzymatic dissociation masks mitochondrial Ca^2+^ handling defects in mitochondrial myopathy muscle fibers

To evaluate the effect of enzymatic dissociation on muscle fiber Ca^2+^ handling in more detail, we used fibers from fast-twitch skeletal muscle fiber-specific mitochondrial transcription factor A knock-out (*Tfam* KO) mice that display important hallmarks of severe mitochondrial myopathy [54]. In agreement with previous results [42, 51], repeated tetanic stimulation resulted in a marked increase in [Ca^2+^]_mit_ in mechanically dissected *Tfam* KO FDB fibers, whereas [Ca^2+^]_mit_ was maintained at a low level in muscle fibers of non-KO littermate controls (Figure 8A). This increased mitochondrial Ca^2+^ uptake in *Tfam* KO fibers occurred despite a reduced driving force for Ca^2+^ into the mitochondrial matrix due to lower tetanic [Ca^2+^]_cyt_ in *Tfam* KO than in control fibers (Figure 8B) [42, 51]. Importantly however, when enzymatic dissociated fibers were used, also the control non-KO fibers showed extensive elevations of [Ca^2+^]_mit_ and any difference in mitochondrial Ca^2+^ uptake between *Tfam* KO and control fibers was lost. These findings corroborate the dystrophic phenotype of enzymatically dissociated fibers and demonstrates that only microdissected fibers can fully reveal disease-specific defects in mitochondrial Ca^2+^ handing.

**Figure 8.**
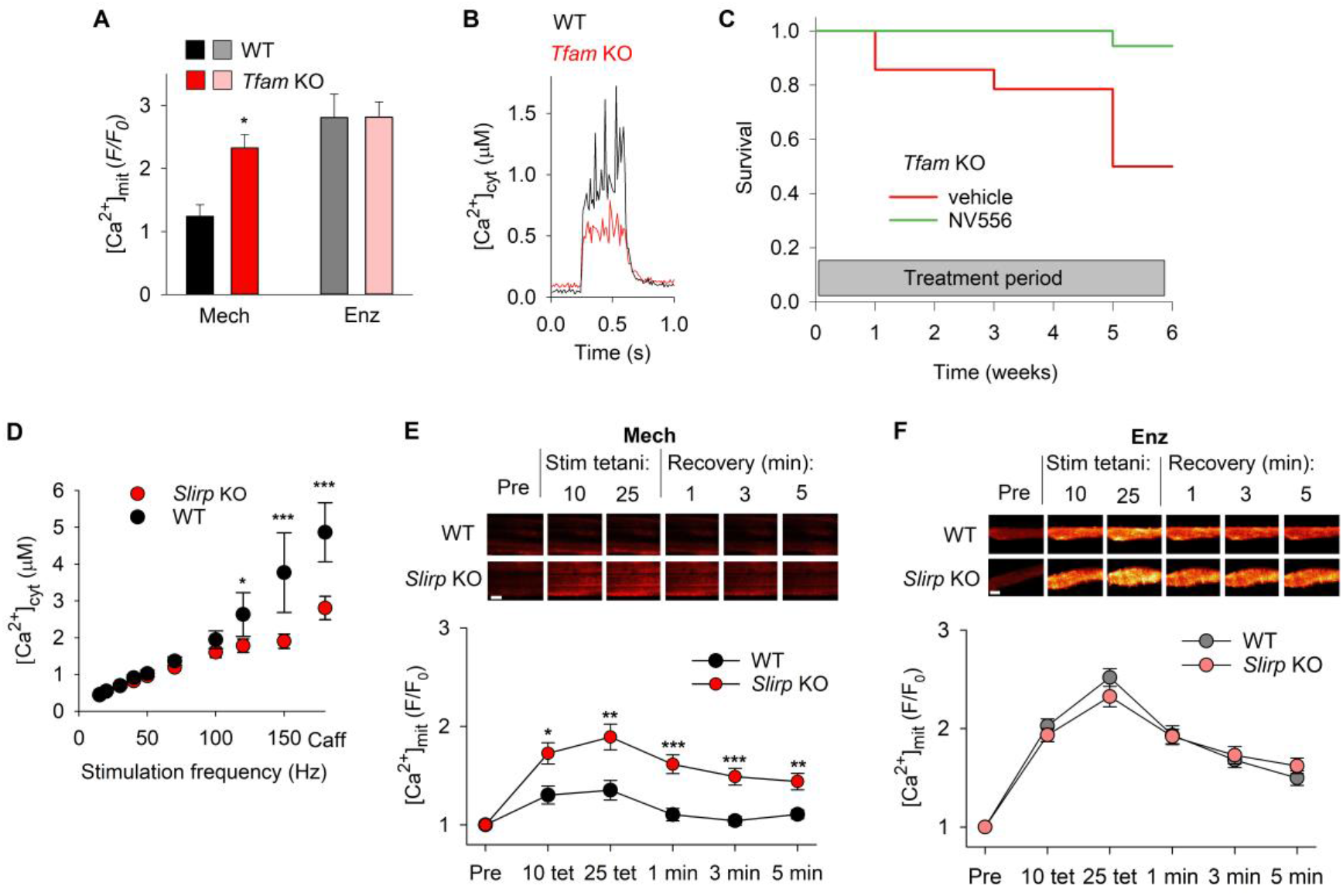
The aberrant contraction-induced mitochondrial Ca^2+^ uptake in mitochondrial myopathy muscle fibers eludes detection in enzymatically dissociated fibers. **(A)** Mean data of the relative increase in rhod-2 fluorescence after 25 repeated tetanic contractions in mechanically dissected (Mech) Tfam KO (n = 5) and littermate controls (WT; n = 3) or enzymatically dissociated (Enz) Tfam KO (n = 10) and WT (n = 4) FDB fibers. * P < 0.05 vs. WT with unpaired t-test. **(B)** Superimposed average [Ca^2+^]_cyt_ records during tetanic stimulation (70 Hz, 350 ms duration) obtained in mechanically dissected FDB fibers of control mice (WT, n = 6 fibers; same data as in Figure 3B) and Tfam KO mice (n = 4 fibers). (**C**) Survival curves Tfam KO mice treated with the PPIF inhibitor NV556 that blocks mitochondrial Ca^2+^ uptake (140 µg daily; green line) or with vehicle control (red line) using osmotic pumps for up to 6 weeks (treatments started at an age of 14 weeks). (**D**) Mean data of cytosolic free [Ca^2+^] ([Ca^2+^]_cyt_) during brief stimulations at different frequencies and in the presence of caffeine (Caff, 5 mM) of WT (n = 7) and Slirp KO (n = 11) FDB fibers. Representative rhod-2 fluorescence images of a small bundle of mechanically dissected (**E**) and enzymatically dissociated (**F**) FDB fibers of WT and Slirp KO mice obtained before (Pre) and after 10 and 25 repeated tetanic stimulation and at 1, 3, and 5 minutes of recovery (scale bars, 20 µm). Lower parts of E and F show mean data of the stimulation-induced relative increase in rhod-2 fluorescence (n ≥ 19). Experiments were performed 4 hours after fiber isolation. * P < 0.05, ** P < 0.01, *** P < 0.001 vs. WT with two-way repeated measures ANOVA. Data are presented as mean ± SEM.

To test whether PPIF inhibition would ameliorate the dystrophic effects, we evaluated survival of *Tfam* KO mice treated with NV556. To this end, NV556 was delivered via osmotic pumps and treatment started at an age of 14 weeks when the *Tfam* KO mice are about to enter the terminal state with severe weight loss and muscle weakness [51]. The osmotic pumps delivered ∼140 µg NV556 per day for up to 6 weeks. At the end of the treatment period, 17 out of 18 NV556-treated mice were still alive, whereas only 7 out of 14 untreated mice were alive (*P* = 0.015; z-test; Figure 8C). These data further support the critical role of PPIF in mitochondrial Ca^2+^ control and show that pharmacological inhibition of excessive mitochondrial Ca^2+^ uptake improves outcomes in a mouse model of lethal mitochondrial myopathy. Intrigued by fact that the aberrant mitochondrial Ca^2+^ uptake in *Tfam* KO fibers eluded detection in enzymatically dissociated fibers, we performed experiments on another mouse model with defective mitochondria; that is, mice deficient of stem-loop interacting RNA binding protein (SLIRP) [55]. Despite a 50–70% reduction in the steady-state levels of mtDNA-encoded mRNAs, *Slirp* KO mice appeared largely healthy with only minor (∼5%) reduction in body weight [55]. The [Ca^2+^]_cyt_-frequency relationship was studied in FDB fibers mechanically dissected from *Slirp* KO mice and wildtype littermates by producing brief contractions at 1 minute intervals. *Slirp* KO fibers displayed lower [Ca^2+^]_cyt_ than wildtype fibers at high stimulation frequencies (120-150 Hz) and during tetanic stimulation in the presence of caffeine (5 mM), which facilitates SR Ca^2+^ release and hence provides an estimate of the total amount of Ca^2+^ stored in the SR (Figure 8D) [56]. These results indicate a reduced SR Ca^2+^ storage capacity in *Slirp* KO muscle, which agrees with previous results obtained in *Tfam* KO fibers [42, 51]. Thus, a decreased SR Ca^2+^ storage capacity is a common feature in muscle fibers of mice with two completely different genetically engineered mitochondrial defects.

In accordance with the results from *Tfam* KO muscle fibers, rhod-2 fluorescence increased more during 25 repeated tetanic contractions in mechanically dissected *Slirp* KO than in wildtype muscle fibers (Figure 8E). Importantly, this difference between *Slirp* KO and wildtype fibers was masked by substantial increases in [Ca^2+^]_mit_ in the wildtype group when experiments were performed with enzymatically dissociated fibers (Figure 8F). Thus, two mitochondrial myopathy mouse models, *Slirp* KO and *Tfam* KO, both showed augmented mitochondrial Ca^2+^ uptake during repeated contractions despite decreased SR Ca^2+^ storage and hence, if anything, decreased [Ca^2+^]_cyt_ during tetanic contractions. Importantly, isolation of wildtype muscle fibers by enzymatic ECM digestion perturbed mitochondrial Ca^2+^ handling such that it resembled the defective Ca^2+^ handling observed in mitochondrial myopathy models. Consequently, differences between control and myopathy fibers were masked in enzymatically dissociated fibers. In contrast, fibers isolated by microdissection with preserved extracellular microenvironments, allowed detection of the aberrant mitochondrial in muscle fibers of both *Tfam* KO and *Slirp* KO mitochondrial myopathy models.

## Discussion

*In vitro* cell studies constitute essential tools for studies of cell biology, morphogenesis, phenotype characterization, as well as for drug development. However, an increasing body of evidence highlights the fact that isolating cells by enzymatic dissociation and subsequently studying them for several days entails the rapid loss of adult cellular phenotypes, which confounds result interpretation and impairs the translation of findings [57]. Enzymatic dissociation of cells disturbs integrin-mediated cell adhesions, which provide dynamic connections between the ECM and the intracellular cytoskeleton that are essential for the control of cell structure, including the morphology and function of mitochondria [19–21]. In this study, we provide a link between enzymatic disruption of the ECM, altered cell structure, and defective control of mitochondrial Ca^2+^ uptake by comparing enzymatically dissociated muscle fibers with muscle fibers isolated by mechanical dissection, which leaves the immediate ECM intact.

Enzymatic dissociation resulted in a loss of structural integrity and drastically reduced expression of genes encoding for structural proteins, such as collagens and matrix proteoglycans. Moreover, FIB-SEM revealed subtle differences in the mitochondrial 3D network with a higher number of mitochondria with individually lower volumes in enzymatically dissociated fibers, hence supporting an important role of the ECM in orchestrating the dynamic balance between mitochondrial fusion and fission [58–60]. In contrast, genes associated with cell cycle and DNA replication were significantly induced in enzymatically dissociated compared to microdissected fibers. These fundamental alterations were paralleled by a downregulation in enzymatically dissociated fibers of markers of skeletal muscle maturation, such as genes for myosin heavy chains. Notably, gene expression of the actin regulator gelsolin was maintained in microdissected fibers, whereas it was among the most downregulated genes in enzymatically dissociated fibers. Gelsolin is inactive in the absence of Ca^2+^; however, upon Ca^2+^ binding, gelsolin undergoes conformational changes, and the resulting activated domains participate in the severing and capping of actin filaments [61–63]. Moreover, gelsolin has been shown to inhibit the cellular stress-induced increase in mitochondrial membrane permeability and loss of mitochondrial membrane potential in a Ca^2+^- and CsA-dependent manner [64], i.e. acting on processes that involve Ppif [47–50]. Combined, these findings argue for a rapid dedifferentiation of adult muscle fibers upon isolation from their native microenvironment, which corroborates with previous results showing that ECM has a central role in the skeletal muscle differentiation process [65]. These effects resemble observations in other tissues, such as liver, brain, and heart [66–69], and argue for the importance of maintaining the physiological niche in *ex vivo* experiments of cellular function.

Our results imply that the aberrant mitochondrial Ca^2+^ uptake in enzymatically dissociated fibers is PPIF-dependent as the excessive increase in [Ca^2+^]_mit_ in response to increased [Ca^2+^]_cyt_ was significantly decreased by the cyclophilin inhibitors CsA and NV556. Notably, the excessive increase in [Ca^2+^]_mit_ in enzymatically dissociated fibers during repeated tetanic stimulation was observed already 4 hours after cell isolation, hence too soon for important changes in protein levels to develop. Thus, the excessive mitochondrial Ca^2+^ uptake would be a direct consequence of altered mitochondrial function caused by disrupted ECM and intracellular cytoskeleton. Nevertheless, changes in gene expression 24 hours after muscle fiber isolation indicate an additional enzymatic dissociation-induced long-term shift towards increased [Ca^2+^]_mit_; that is, the expression of *Mcu* and *Ppif* was higher and the expression of *Nclx* was lower in enzymatically dissociated than in microdissected fibers, which would promote mitochondrial Ca^2+^ influx and limit Ca^2+^ extrusion. Interestingly, previous studies showed that myopathies in mice deficient in the ECM protein collagen VI could be counteracted by CsA [30] and the non-immunosuppressive Ppif inhibitor alisporivir (also called Debio 025) [31]. Furthermore, our finding that pharmacological inhibition of aberrant mitochondrial Ca^2+^ uptake improved survival in the *Tfam* KO mouse model of lethal mitochondrial myopathy supports a scheme where the adverse effects of ECM perturbations are mediated, at least in part, via impaired Ppif-dependent fine-tuning of cytosolic-mitochondrial Ca^2+^ fluxes, thus providing a molecular explanation for prolonged survival previously reported when *Tfam* KO mice were treated with CsA [51], as well as treatment with the more specific cyclophilin inhibitor NV556 used in the present study.

The presented findings highlight multiple important implications for the study of cell biological phenomena *in vitro*. Firstly, our data emphasize the importance of using cell culture systems that preserve the immediate ECM to ensure that results faithfully reflect *in vivo* processes. Secondly, enzymatic digestion rapidly changes the molecular phenotypes and functionality of cells. For instance, recent studies have shown redundant activation of muscle stem cells isolated from adult skeletal muscle with standard enzymatic dissociation protocols, which has important consequences for the use of these cells as quiescent controls [70, 71]. Thirdly, dissolving the microphysiological niche around cells can result in perturbations that resemble pathological phenotypes observed in mitochondrial disease, providing further evidence for an intricate interplay between cellular structure, Ca^2+^ fluxes, metabolism, and function. Specifically, the pathognomonic increase in [Ca^2+^]_mit_ during repeated contractions of muscle fibers in mitochondrial myopathy would be missed in experiments performed on enzymatically dissociated cells. Thus, enzymatically dissociated cells should be avoided as an experimental paradigm for the study of diseases that potentially involve altered mitochondrial Ca^2+^ signaling.

In conclusion, disruption of the organotypic niche results in the loss of structural integrity of muscle fibers accompanied by impaired control of mitochondrial Ca^2+^ (Figure 9). The molecular link between the processes involves a Ppif-dependent mitochondrial Ca^2+^ influx resembling that observed in mitochondrial myopathies. Our results support a central role of mitochondrial Ca^2+^ as a critical mediator that connects the native extracellular microenvironment to the maintenance of normal cellular structure and function.

**Figure 9.**
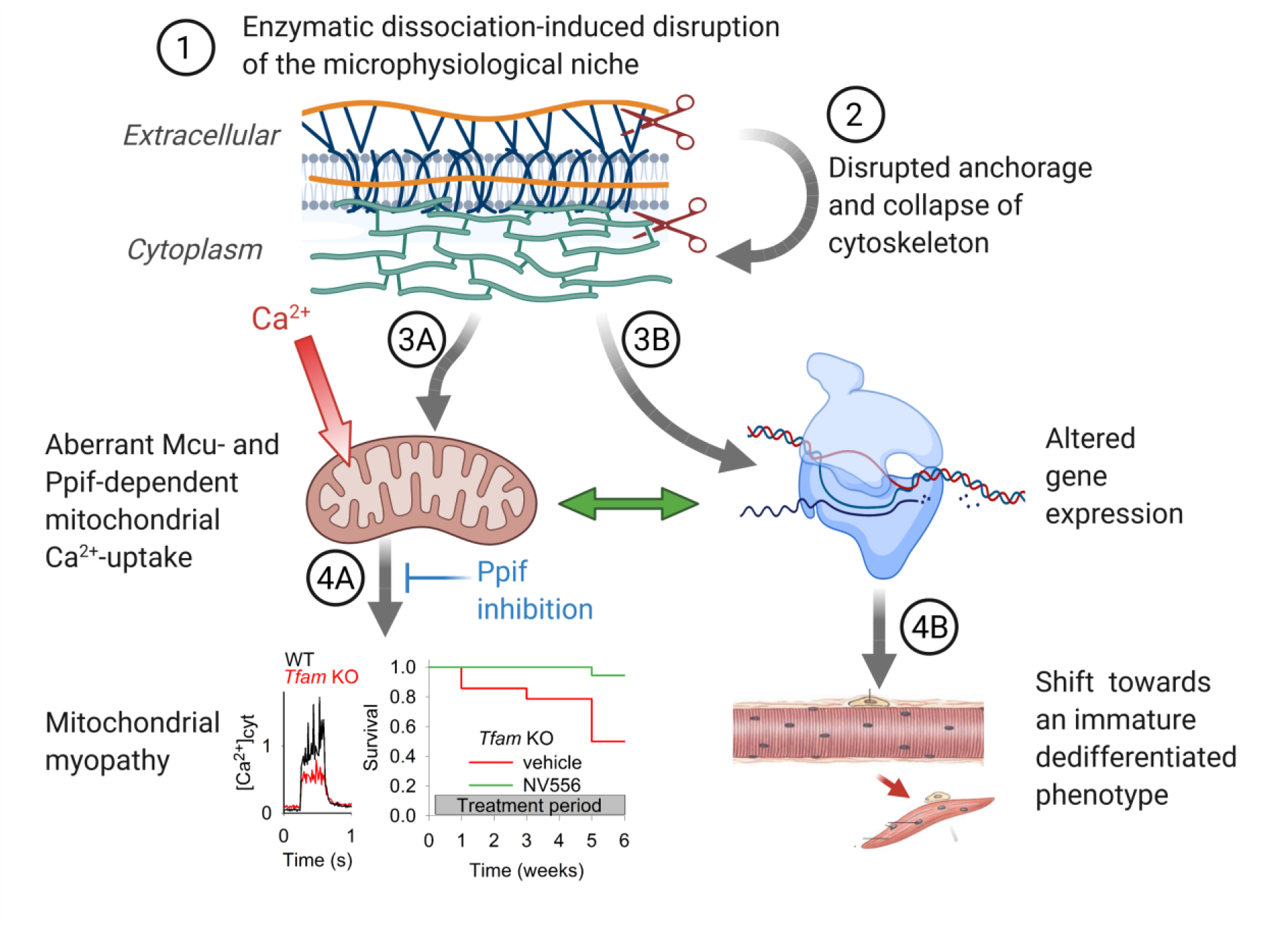
Schematic summary of major results. Muscle fiber isolation by enzymatic dissociation disrupts the ECM (**1**), which results in collapse of the cytoskeleton and disruption of the fine-detailed muscle fiber structure (**2**). This in turn results in aberrant mitochondrial Ca^2+^ accumulation, which is partly blocked by inhibition of either Mcu or Ppif (**3A**). Pharmacological Ppif inhibition of the aberrant mitochondrial Ca^2+^ uptake counteracts the development of muscle weakness and prolongs the lifespan of Tfam KO mitochondrial myopathy mice (**4A**), and a similar mechanism might also underlie the positive effects of Ppif inhibition in collagen VI mutation-dependent muscular dystrophies [28–31]. Enzymatic dissociation also shifts the pattern of gene expression (**3B**), resulting in a shift towards an immature dedifferentiated phenotype (**4B**). Several probable interactions between the A- and B-arm of ECM disruption-induced impairments are revealed (green doubleheaded arrow), but the precise functional interactions between the two arms remain to be defined. Created with BioRender.com.

## Materials and methods

### Animals

C57bl/6J female mice were used, except when stated otherwise. Fast-twitch skeletal muscle fiber-specific *Tfam* KO mice and their controls were generated as described previously [54]. *Slirp* KO mice and wildtype littermates were generated as described previously [55]. Mice were euthanized by rapid neck dislocation and the FDB muscles were excised. All animal experiments were approved by the Stockholm North Local Animal Ethics Committee.

Some *Tfam* KO mice were treated with the specific cyclophilin inhibitor, NV556 [53], or vehicle only. NV556 was dissolved to 40 mg/ml in cyclodextrin formulation (pH 7.4). Mini-osmotic pumps (Alzet model 2006) were loaded with approximately 200 µl of either dissolved NV556 or cyclodextrin formulation only and incubated in sterile PBS for 48 hours prior to implantation. The mini-osmotic pumps were implanted subcutaneously on the back of mice under isoflurane anaesthesia. Mice were weighed 48 hours after surgery; body weight was then monitored twice per week for the remainder of the experiment period. Mice were regarded as terminally ill and euthanized either at loss of 20% body weight or severe loss of muscle functionality. Mice were euthanized by rapid neck dislocation.

### Enzymatic dissociation and mechanical dissection of FDB fibers

*Flexor digitorum brevis* (FDB) muscles were cleaned of tendons, connective tissue, and blood vessels and incubated for ∼2 hours at 37°C in 0.3% collagenase type I (Sigma) in Dulbecco’s modified Eagle medium (DMEM; Invitrogen) supplemented with 20% fetal bovine serum. Muscles were then transferred to fresh DMEM and gently triturated to dissociate individual muscle fibers. A volume of 300 μl of the resultant muscle fiber suspension was placed in laminin-coated glass-bottom Petri dishes, and fibers were allowed to attach for up to 15 minutes. Thereafter, 3 ml DMEM supplemented with antibiotic, antimycotic solution (1 μL/ml, Sigma) was added. Experiments were performed ∼4 hours after enzymatic dissociation, except when otherwise noted.

Single or small bundles (2-5 fibers) of FDB fibers were mechanically dissected as previously described [37]. In one set of experiments (see Figure 5C), dissected fibers were subjected to the same enzymatic dissociation protocol as the dissociated fibers with the exception that no trituration was performed.

### Second Harmonic Generation microscopy

FDB fibers were imaged using a multiphoton microscope (TriMScope II, LaVision BioTech, Bielefeld, Germany) equipped with a combination of two water immersion objectives (LD C-APO 40x/1.1 W Corr M27 on the excitation side, W Plan-APO 20x/1.0 DIC M27 on the transmission side, Zeiss) and a mode-locked ps-pulsed Ti:Sa-laser (Chameleon Vision II, Coherent, Santa Clara, CA) tuned to 810 nm to excite the myosin SHG signal [72, 73]. The SHG signal from myosin was separated from other (autofluorescence) signals by a band pass filter (405/20 nm, CHROMA, Bellows Falls, VT) and detected by a non-descanned photomultiplier (H7422-40, Hamamatsu Photonics, Hamamatsu, Japan). Muscle fibers were z-scanned (voxel-size: 0.14 × 0.14 × 0.3 µm) to detect the cosine angle sum based on boundary tensor analysis [38, 39]. Image processing was performed in MATLAB (MathWorks, Natick, MA).

### Transmission Electron Microscopy

Isolated FDB muscle fibers were fixed in 2.5% glutaraldehyde (TAAB Laboratories), 4% formaldehyde in 0.1 M sodium cacodylate buffer. All samples were processed using Pelco Biowave Pro+ microwave tissue processor (Ted Pella) according to [74], with minor modifications: no Ca^2+^ was used during the fixation and contrasting steps with lead aspartate were omitted to reduce overstaining. Samples were trimmed with a glass knife and 70 nm ultrathin sections were picked up on Cu-grids and examined with the TEM Talos L120C (FEI, currently Thermo Fischer Scientific) operating at 120 kV. Micrographs were acquired with a Ceta 16M CCD camera (FEI) using TEM Image & Analysis software ver. 4.17 (FEI).

For focused ion beam scanning electron microscopy (FIB-SEM), a small cube of the sample was cut and glued on an SEM stub with epoxy and silver glue. In the SEM chamber, the specimen was coated with a 5 nm thin layer of platinum to reduce charging. Specimens were imaged using Scios DualBeam SEM (FEI) and the ‘Auto slice and view’ software system; the electron beam operated at 2 kV and 0.2 nA and was detected with T1 in-lens detector. A 1 µm protective layer of platinum was deposited on the selected area before milling. FIB milling thickness was set to 30 nm and each slice was imaged with pixel sizes 3.55 × 3.55 nm (for mechanically dissected fiber) and 3.74 × 3.74 nm (for enzymatically dissociated fiber). Images were further processed using the ImageJ plugins ‘Linear Stack Alignment with SIFT’ and ‘Multistackreg’ and the mitochondrial network was reconstructed and analyzed in a final volume of 365 µm^3^ (7.9 × 4.5 × 10.3 µm). Identified mitochondrial volumes were modelled and measured using the IMOD software package ver. 4.9.13 [75].

### RNA-Sequencing

RNA sequencing by poly-A capture was performed using >10 ng RNA input material. Genes with an average number of fragments per kilo base per million mapped reads (FPKM) >0.5 across all samples were analyzed using Qlucore (Lund, Sweden). Differential gene expression analysis was conducted using DESeq2 and multiple testing correction was applied using the Benjamini–Hochberg procedure with false discovery rates (FDRs) ≤ 5%. Pathway enrichment analysis was conducted based on the PANTHER gene family classification system using the WebGestalt toolbox [76].

### Confocal measurements with fluorescent indicators in FDB fibers

For mitochondrial [Ca^2+^] ([Ca^2+^]_mit_), membrane potential (Δ*ψ*_m_) and ROS production measurements (see below), loading of fluorescent indicators followed by wash-out was performed at room temperature for 20 minutes. During FDB fiber experiments, cells were superfused at room temperature (∼25°C) with Tyrode solution (in mM): NaCl, 121; KCl, 5.0; CaCl_2_, 1.8; MgCl_2_, 0.5; NaH_2_PO_4_, 0.4; NaHCO_3_, 24.0; EDTA, 0.1; glucose, 5.5; 0.2% fetal calf serum. The solution was bubbled with 95% O_2_–5% CO_2_. Measurements were performed using a BioRad MRC 1024 confocal unit with a dual Calypso laser (Cobolt, Solna, Sweden) attached to a Nikon Diaphot 200 inverted microscope. Confocal images were obtained before and after 10 and 25 repeated tetani (70 Hz, 350 ms stimulation trains given at 2 s intervals) and at regular intervals after the contractions. Confocal images were analyzed using ImageJ and data are expressed as *F*/*F*_0_, i.e. the ratio of the fluorescence intensity after and before the repeated contractions, respectively.

#### [Ca^2+^]_mit_

Fibers were incubated in 5 µM rhod-2 in the membrane permeable AM form (Invitrogen). Rhod-2 was excited with 531 nm light and the emitted light was collected through a 585 nm long-pass filter.

#### Δ*ψ*_m_

Fibers were incubated in 1 µM TMRE (Invitrogen). TMRE was excited at 531 nm and the emitted light was collected through a 605 nm long-pass filter. Approximately 30 minutes after completion of the 25 repeated tetani, fibers were exposed to 1 µM FCCP (Sigma) to significantly depolarize the mitochondria and confocal images were obtained.

#### Mitochondrial ROS production

Fibers were loaded with 5 µM MitoSOX Red (Invitrogen). MitoSOX Red was excited with 488 nm light and emitted light was collected through a 585 nm long-pass filter. As a positive control, 1 mM H_2_O_2_ was applied at the end of experiments to increase mitochondrial superoxide by inducing product inhibition of superoxide dismutase 2 [77, 78], and confocal images were obtained every minute until the MitoSOX Red signal reached a plateau.

#### Ru360, CsA and NV556 experiments

Enzymatically dissociated FDB fibers were loaded with rhod-2 to measure [Ca^2+^]_mit_ as described above. To investigate the potential sites of Ca^2+^ entry into mitochondria, enzymatically dissociated fibers were exposed to either Ru360 (Calbiochem), CsA or NV556 (NeuroVive Pharmaceutical AB, Lund, Sweden). Ru360 was first injected into the fibers and they were subsequently superfused with 10 µM Ru360 throughout the experiment. CsA (1.6 μM; Novartis) or NV556 (5 µM) was applied to the fibers for 5 minutes before and during the 25 repeated tetani.

### Single fiber [Ca^2+^]_cyt_ measurements

Intact single FDB fibers were mechanically dissected [37]. Aluminum or platinum clips were attached to the tendons and the fiber was mounted in a chamber between an Akers 801 force transducer (Kronex Technologies, Oakland, CA, USA) and an adjustable holder and subsequently superfused by Tyrode solution (see above) at room temperature. The fiber length was adjusted to obtain maximum tetanic force. The fiber was stimulated with supramaximal electrical pulses (0.5 ms in duration) delivered via platinum electrodes placed along the long axis of the fiber. The steady-state [Ca^2+^]_cyt_-frequency relationship was obtained by stimulating fibers for 350 ms at 15-150 Hz every 1 minutes; 150 Hz contractions were also produced in the presence of 5 mM caffeine to assess the SR Ca^2+^ storage capacity.

Fibers were microinjected with the fluorescent Ca^2+^ indicator indo-1 (Molecular Probes/Invitrogen, Carlsbad, CA, USA), or loaded with indo-1 AM in experiments on enzymatically dissociated fibers. The emitted fluorescence of indo-1 was measured with a system consisting of a Xenon lamp, a monochromator, and two photomultiplier tubes (Photon Technology International, Wedel, Germany). The excitation light was set to 360 nm, and the light emitted at 405 ± 5 and 495 ± 5 nm was measured by the photomultipliers. The ratio of the light emitted at 405 nm to that at 495 nm (R) was converted to [Ca^2+^]_cyt_ using the following equation:

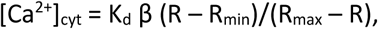

where K_d_ is the apparent dissociation constant of indo 1, β is the ratio of the 495 nm signals at very low and saturating [Ca^2+^]_cyt_, R_min_ is the ratio at very low [Ca^2+^]_cyt_, and R_max_ is the ratio at saturating [Ca^2+^]_cyt_. Fluorescence was sampled online and stored on a computer for subsequent data analysis. [Ca^2+^]_cyt_ was measured at rest and as the mean during 70 Hz, 350 ms tetanic stimulation. The steady-state [Ca^2+^]_cyt_-frequency relationship was obtained in *Slirp* KO and control WT fibers by stimulating fibers for 350 ms at 15-150 Hz every 1 minutes; 150 Hz contractions were also produced in the presence of 5 mM caffeine to assess the SR Ca^2+^ storage capacity.

### Statistics

Statistical analyses were performed with SigmaPlot 13 (Systat Software Inc.). Student’s paired, unpaired t-tests, one-way ANOVA, or z-test were used as appropriate. Two-way repeated measures ANOVA was used to determine differences between two groups of repeatedly stimulated fibers. The Holm-Sidak post-hoc analysis was used when ANOVA showed a significant difference between groups. Significance was assumed for *P* < 0.05. Data are presented as mean ± SEM.

## Supporting information

Video 1 SHG mechanical

Video 2 SHG enzymatic

Video 3 EM mechanical 3D mitochondria

Video 4 EM enzymatic 3D mitochondria

## Acknowledgements

This work was supported by grants from the Swedish Research Council Swedish Research Council (2018‐02576), the Swedish Research Council for Sport Science (P2019‐0060), Association Française contre les Myopathies (AFM-Téléthon #16798; to C.G.), and the German Research Foundation DFG (FR2993/13-1). For technical support and EM access, we acknowledge the Umeå Centre for Electron Microscopy, the SciLifeLab at Umeå University, the National Microscopy Infrastructure (NMI), including instruments funded by the Knut and Alice Wallenberg foundation and Kempe foundations. Research reported in this publication was supported by the National Institute of Arthritis and Musculoskeletal and Skin Diseases of the National Institutes of Health under award number 1F32AR057619 (NIAMS; to A.H.). The content is solely the responsibility of the authors and does not necessarily represent the official views of the National Institutes of Health.

## Author Contributions

C.G., S.U.V., S.Y, A.H., J.D.B., N.I., V.M.L. and H.W. conceived the project and designed the experiments. C.G., S.U.V., S.Y., A.H., A.J.C., T.C., J.D.B. and N.I. performed most of the experiments. A.B., D.S. and O.F. performed Second Harmonic Generation microscopy experiments. K.H. performed TEM experiments. S.H. and L.S. performed FIB-SEM experiments. C.G., N.I., V.M.L. and H.W. wrote the manuscript. All authors contributed to the editing of the manuscript.

