## Supplementary figures for "Disruption of the microphysiological niche alters matrix deposition and causes loss of mitochondrial Ca^2+^ control in skeletal muscle fibers"

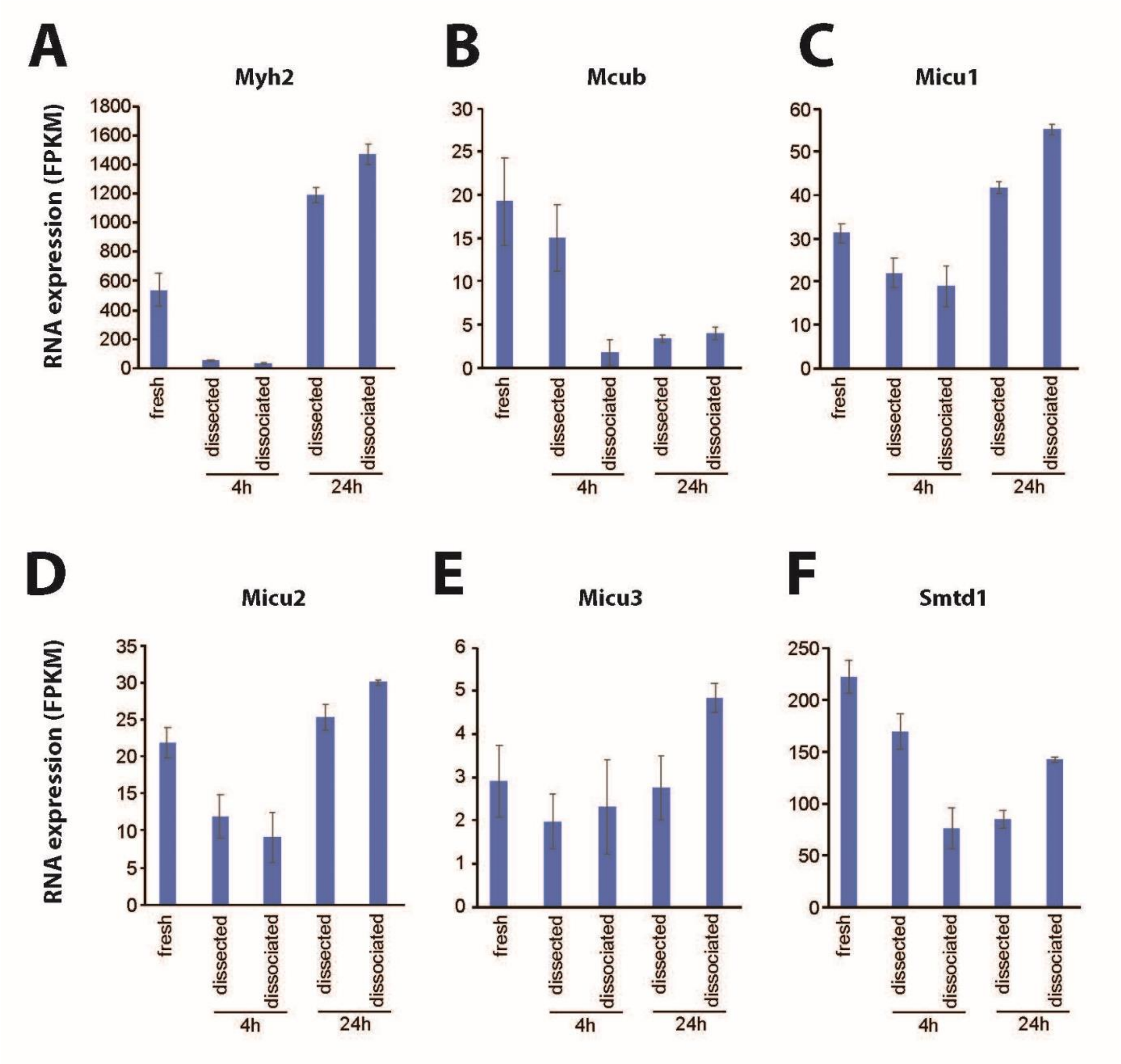


**Supplementary Figure 1. Enzymatic dissociation had no consistent effect on the mRNA expression of myosin heavy chain IIA or Mcu regulators.** Expression of genes encoding for myosin heavy chain IIA (Myh2; **A**) and Mcu regulators: mitochondrial $Ca^{2+}$ uniporter B (Mcub; **B**); mitochondrial $Ca^{2+}$ uptake proteins 1-3 (Micu1-3; **C-E**); and essential Mcu regulator (EMRE, Smtd1; **F**). Data are presented as mean ± SEM; n=4.

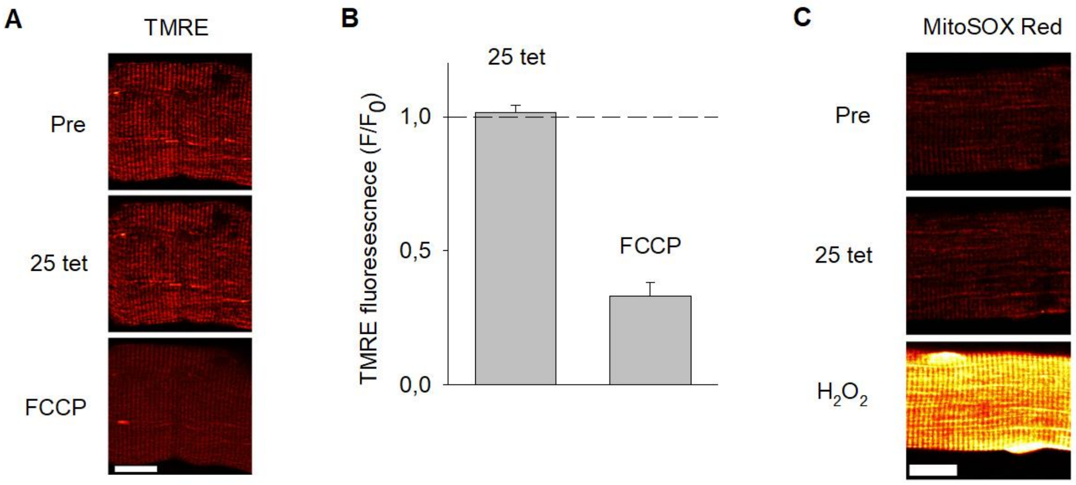


**Supplementary Figure 2. Enzymatic dissociation does not cause mitochondrial depolarization or increased ROS production.**

(**A**) Representative TMRE fluorescence images and (**B**) mean data (± SEM) show no significant increase in TMRE fluorescence after (F) relative to before ($F_0$) 25 repeated tetani, whereas the subsequent exposure to the mitochondrial uncoupler FCCP (1 µM) resulted in a marked decrease in fluorescence (n = 17). (**C**) Representative images show no clear increase in MitoSOX Red fluorescence after 25 contractions, whereas fluorescence increased several-fold during the subsequent exposure to 1 mM $H_2O_2$. Experiments were performed 4 hours after fiber isolation. Scale bars in **A** and **C**, 20 µm.
